# Genomic Influences on Self-Reported Childhood Maltreatment

**DOI:** 10.1101/717314

**Authors:** Shareefa Dalvie, Adam X. Maihofer, Jonathan R.I. Coleman, Bekh Bradley, Gerome Breen, Leslie A. Brick, Chia-Yen Chen, Karmel W. Choi, Laramie E. Duncan, Guia Guffanti, Magali Haas, Supriya Harnal, Israel Liberzon, Nicole R. Nugent, Allison C. Provost, Kerry J. Ressler, Katy Torres, Ananda B. Amstadter, S. Bryn Austin, Dewleen G. Baker, Elizabeth A. Bolger, Richard A. Bryant, Joseph R. Calabrese, Douglas L. Delahanty, Lindsay A. Farrer, Norah C. Feeny, Janine D. Flory, David Forbes, Sandro Galea, Aarti Gautam, Joel Gelernter, Rasha Hammamieh, Marti Jett, Angela G. Junglen, Milissa L. Kaufman, Ronald C. Kessler, Alaptagin Khan, Henry R. Kranzler, Lauren A. M. Lebois, Charles Marmar, Matig R. Mavissakalian, Alexander McFarlane, Meaghan O’Donnell, Holly K. Orcutt, Robert H. Pietrzak, Victoria B. Risbrough, Andrea L. Roberts, Alex O. Rothbaum, P. Roy-Byrne, Ken Ruggiero, Antonia V. Seligowski, Christina M. Sheerin, Derrick Silove, Jordan W. Smoller, Nadia Solovieff, Murray B. Stein, Martin H. Teicher, Robert J. Ursano, Miranda Van Hooff, Sherry Winternitz, Jonathan D. Wolff, Rachel Yehuda, Hongyu Zhao, Lori A. Zoellner, Dan J. Stein, Karestan C. Koenen, Caroline M. Nievergelt

## Abstract

Childhood maltreatment is highly prevalent and serves as a risk factor for mental and physical disorders. Self-reported childhood maltreatment appears heritable, but the specific genetic influences on this phenotype are largely unknown. The aims of this study were to 1) identify genetic variation associated with reported childhood maltreatment, 2) calculate the relevant SNP-based heritability estimates, and 3) quantify the genetic overlap of reported childhood maltreatment with mental and physical health-related phenotypes. Genome-wide association analysis for childhood maltreatment was undertaken, using a discovery sample from the UK Biobank (UKBB) (n=124,000) and a replication sample from the Psychiatric Genomics Consortium–posttraumatic stress disorder working group (PGC-PTSD) (n=26,290). Heritability estimations for childhood maltreatment and genetic correlations with mental/physical health traits were calculated using linkage disequilibrium score regression (LDSR). Two genome-wide significant loci associated with childhood maltreatment, located on chromosomes 3p13 (rs142346759, beta=0.015, p=4.35×10^−8^, *FOXP1*) and 7q31.1 (rs10262462, beta=-0.016, p=3.24×10^−8^, *FOXP2*), were identified in the discovery dataset but were not replicated in the PGC-PTSD sample. SNP-based heritability for childhood maltreatment was estimated to be ∼6%. Childhood maltreatment was most significantly genetically correlated with depressive symptoms (r_g_=0.70, p=4.65×10^−40^). This is the first large-scale genetic study to identify specific variants associated with self-reported childhood maltreatment. *FOXP* genes could influence traits such as depression and thereby be relevant to childhood maltreatment. Alternatively, these variants may be associated with a greater likelihood of reporting maltreatment. A clearer understanding of the genetic relationships of childhood maltreatment, including particular abuse subtypes, with various psychiatric disorders, may ultimately be useful in in developing targeted treatment and prevention strategies.

## Introduction

The lifetime prevalence of childhood physical, sexual and emotional abuse ranges from 8-36%^1^. In addition to being highly prevalent, childhood maltreatment is associated with the development of mental disorders, including depression^2,3^, and physical ill health, including non-communicable diseases^4,5^. Although these associations are now well established, estimates of effect size vary considerably across epidemiological studies, likely reflecting methodological challenges, including uncertainty about how best to assess childhood maltreatment ^6^.

A twin-based study found that retrospective reports of childhood maltreatment has a heritability of 6% ^7^. Although the idea that childhood maltreatment is heritable may be counter-intuitive, the field of behavior genetics has long documented the heritability of many exposures seen as environmental. This is referred to as gene-environment correlation (rGE). Three potential rGE mechanisms to explain the heritability of childhood maltreatment may be posited. First, a “passive” rGE: parental genes affecting parental behavior may influence the childhood environment (e.g. aggressive parents may be more likely to physically punish their children who have also inherited their parents’ genetic variants influencing aggressive behavior ^8^). Second, an “active” rGE: individuals with genetic variants associated with certain behavioral phenotypes may be more at risk of selecting or creating adverse situations (e.g. risk-taking personality is heritable and children who are high in risk taking may be exposed to more trauma)^9,10^. Third, an “evocative” rGE: genetic variation may influence child behavior, which in turn is associated with responses to the child (e.g. genetic factors may influence infant “difficultness”, which in turn is associated with maternal hostile-reactive behavior that is correlated with child abuse^11,12^). The latter two rGEs are sometimes collectively referred to as non-passive correlations^7^.

While a number of key risk factors for childhood maltreatment, including child behavioral characteristics and parental mental health, have been investigated^6^, studies have seldom focused on associated genetic variation. The few genetic association studies of childhood maltreatment have only considered variants in candidate genes^13^ and have had insufficient power to detect the small polygenic effect sizes typically associated with behavioral phenotypes^14^. Also, there are no studies of the genetic overlap of childhood maltreatment with mental and physical health-related traits, using genome-wide single nucleotide polymorphism (SNP) data. Knowledge of specific genetic variation for childhood maltreatment, the heritability of this phenotype, and the genetic overlap with other traits may be useful in informing our understanding of the identified risk factors, the etiology and the outcomes of childhood maltreatment. This, in turn, may have important implications for the design of prevention and treatment programs for adverse health outcomes. For example, an observed association between an adverse health outcome and genetic variants may be mediated by an environmental exposure, if that exposure is shown to be causal of the health outcome. Thus, preventative strategies would focus on decreasing the risk conferred by the environmental exposure without needing to specifically consider the genetic influences on the health outcome^9^.

The PGC-PTSD consortium has collaborated to obtain access to well-powered genetic studies of trauma and PTSD that have allowed a number of key genetic questions in this field to be investigated^15-17^, providing a unique opportunity to address knowledge gaps in the area of childhood maltreatment. This study aims to: 1) identify genetic variants associated with childhood maltreatment using a genome-wide association study (GWAS) design, 2) quantify the heritability of childhood maltreatment using SNP-based methods, and 3) assess the degree of genetic overlap of childhood maltreatment with mental and physical health-related phenotypes.

## Methods

### Participating studies

Nineteen studies, comprising subjects of European ancestry only, were used in this analysis. The discovery dataset consisted of 124,711 individuals from the UK Biobank (UKBB)^18^ and the replication sample comprised 26,290 individuals, a subset of the PGC-PTSD Freeze 1.5 dataset (PGC1.5)^17^. The details of these studies, including the demographics and instruments used to assess maltreatment can be found in **Supplementary Table 1**.

### Phenotype Harmonization

For the childhood maltreatment phenotype, Childhood Trauma Questionnaire (CTQ) scores on physical, sexual, and emotional abuse ^19^ were obtained from the participating studies. From this, an overall childhood maltreatment count score of 0-3 was constructed, based on a count of the three abuse categories listed above. An individual was considered to have endorsed a childhood abuse category if they scored in the moderate to extreme range for that particular category, per established cut-offs ^20^ (**Supplementary Table 2**). If CTQ data were not available, the event assessment during childhood (occurring before 18 years of age) that was most validated for that particular study was obtained, providing a count of the total number of different categories of reported childhood events (e.g. physical, sexual or severe emotional abuse) along with the range of possible scores for the measure. The reported maltreatment exposure from the UKBB dataset comprised a score of three items where participants were asked whether they were i) “physically abused by family as a child”, ii) “sexually molested as a child”, and whether they iii) “felt hated by family member as a child”. The childhood maltreatment count score, whether it was generated from the CTQ or another instrument, was used as the main outcome measure in the association analysis. The range of this maltreatment count score for each study can be seen in **Supplementary Table 1**.

### Global Ancestry Determination, Genotyping quality control and Imputation

Study participants from the PGC-PTSD were genotyped with a number of different arrays (**Supplementary Table 1**). Genotype data were quality controlled and processed using the standard PGC pipeline, Ricopili-MANC (https://sites.google.com/a/broadinstitute.org/ricopili/ and https://github.com/orgs/Nealelab/teams/ricopili) as part of the PGC-PTSD Freeze 2 data analysis^17,21^. This work was carried out on the Dutch national e-infrastructure with the support of SURF Cooperative. A detailed outline of these methods can be found in^17^. Briefly, ancestry was determined with pre-QC genotypes using a SNPweights panel of 10,000 ancestry informative markers from a reference panel comprising 2911 subjects from 71 diverse populations and six continental groups (https://github.com/nievergeltlab/global_ancestry). Samples were excluded if they had call rates <98%, deviated from the expected inbreeding coefficient (fhet < −0.2 or > 0.2), or had a sex discrepancy between reported and genotypic sex (based on inbreeding coefficients calculated from SNPs on the X chromosome). Markers were excluded if they had call rates <98%, >2% difference in missing genotypes between PTSD cases and controls, or were monomorphic. Markers with a Hardy-Weinberg equilibrium (HWE) p<1×10^−6^ in controls were excluded from all subjects. Principal components (PCs) were calculated using the smartPCA algorithm in EIGENSTRAT^22^. Pre-phasing and phasing was performed using SHAPEIT2 v2.r837^23^. Imputation was performed with IMPUTE2 v2.2.2^24^ using the 1000 Genomes (1000G) phase 3 data^25^ as the reference.

Details regarding the QC, imputation and ancestry determination of the UKBB dataset can be found in^26^. Briefly, study participants were genotyped with two custom genotyping arrays (with ∼800,000 markers). A two-stage imputation was performed using the Haplotype Reference Consortium (HRC)^27^ and the UK10K^28^ as the reference panels. Variants were filtered to include only those with a minor allele frequency (MAF) of > 1% and an INFO score of > 0.4. Related individuals (third degree and closer) and those with a genotyping call rate < 98% were excluded. Ancestry was determined by 4-means clustering on the first two PCs provided by the UKBB^29^. Additional principal component analysis was conducted on the European-only data subset using flashpca2^30^.

### Main GWAS

GWAS analysis was conducted separately for each study. Best-guess genotypes were tested for association to reported childhood maltreatment using an ordinal logistic regression model with age, sex, and the first five PCs included as covariates. Variants with a MAF < 0.5% and a genotyping rate < 98% were excluded. These analyses were implemented in PLINK 1.9^31^ using the plug-in *Rserve*. To ensure computational efficiency, linear regression models were run for four of the larger contributing studies (NSS1; NSS2; PPDS; and UKBB, N=143,392 subjects)^17^. For the NSS1, NSS2, and PPDS studies, age, sex, and 5 PCs were included as covariates in the regression model. For the UKBB dataset, the regression analysis was implemented in BGenie v1.2^32^ with age, sex, 6 PCs, batch, and site included as covariates.

### Meta-analysis

As both linear and ordinal logistic models were implemented in the GWASs, which resulted in different effect statistics, fixed effects meta-analysis was conducted across studies using p-values and direction of effect, weighted according to the effective sample size as the analysis scheme, in METAL (v. March 25 2011)^33^. Heterogeneity across datasets was tested using the Cochran’s Q-test for heterogeneity, also implemented in METAL. Only variants with an INFO score of greater than 0.8 and a MAF of greater than 5% were included in the meta-analysis, except where otherwise indicated in the results. Forest plots were generated for genome-wide significant hits using the R package *meta*^34^.

### Functional mapping and annotation

Genome-wide significant hits identified from the GWAS meta-analysis were annotated using the web-based tool FUnctional Mapping and Annotation (FUMA) v1.3.4c (http://fuma.ctglab.nl/)^35^. Default settings were used and annotations were based on the human genome assembly GRCh37 (hg19). The *SNP2GENE* module was used to identify genomic risk loci and these were mapped to protein coding genes within a 10kb window. An r^2^ of ≥ 0.6 was used to identify variants in LD with lead SNPs. The 1000G European Phase 3 was used as the reference dataset. Variants were functionally annotated using ANNOVAR, Combined Dependent Depletion (CADD), RegulomeDB (RDB) and chromatin states (only tissues/cells from brains were included). The NHGRI-EBI GWAS catalog was used to determine any previous associations with the identified risk variants. The GTEx v7 brain tissue, RNAseq data from the CommonMind Consortium and the BRAINEAC database were used to perform eQTL mapping for significant SNP-gene pairs (FDR q < 0.05).

A gene-based analysis was performed within FUMA using MAGMA whereby SNPs were mapped to 18,989 protein coding genes. Genome-wide significance was set at a Bonferroni-corrected threshold p<2.63×10^−6^. In addition, gene-based test statistics were used to determine whether specific biological pathways are associated with childhood maltreatment. This was performed for 18,989 curated gene sets and GO terms obtained from MsigDB, using MAGMA. The significance threshold was set at a Bonferroni-corrected threshold of p=2.63×10^−6^ (0.05/18,989).

### Heritability and Genetic Correlation Estimation

Linkage disequilibrium score regression (LDSR) is a technique for quantifying polygenicity and confounding, such as population stratification, in GWAS summary statistics^36^. This is accomplished by evaluating the relationship between linkage disequilibrium scores (the average squared correlation of a SNP with all neighboring SNPs) and SNP test statistics. Using this approach, the LDSR intercept was used to estimate the proportion of inflation in test statistics due to polygenic signal (rather than inflation due to population stratification and cryptic relatedness), with the equation *1 - (LDSR intercept −1)/(mean observed chi-square - 1)*^17^. One of the applications of LDSR is the estimation of SNP-based heritability based on GWAS summary statistics. Another application of LDSR is the measurement of genetic correlation i.e. the degree and direction of shared genetic effects between different traits^36,37^. Heritability and cross-cohort genetic correlation (r_g_) was calculated using LDSR. The web-based interface for LDSR, LD Hub (http://ldsc.broadinstitute.org/ldhub/), was used to further calculate pairwise genetic correlations between childhood maltreatment and 247 non-UKBB traits of interest including psychiatric, anthropomorphic, smoking behavior, reproductive, aging, education, autoimmune, and cardio-metabolic categories.

## Results

### GWAS and meta-analysis

We report GWAS results from our discovery dataset (UKBB) (n=124,711) and meta-analysis (n=151,001). In our UKBB discovery dataset, we identified two genome-wide significant loci (p<5×10^−8^) associated with childhood maltreatment (**Table 1, Figure 1**), rs142346759 (chr3, beta=0.015, p=4.35×10^−8^) and rs10262462 (chr7, beta=-0.016, p=3.24×10^−8^). These variants remained significant in the meta-analysis (**Table 1, Supplementary Figures 1 and 2**). Additional variants on chromosome 7 (rs1859100, beta=0.015, p=3.91×10^−8^) and chromosome 12 (rs917577, beta=0.017, p=2.64×10^−8^) (**Figure 2**), also achieved genome-wide significance in the meta-analysis. None of these hits were replicated in PGC1.5 (**Table 1, Supplementary Figure 3**).

**Figure 1:**
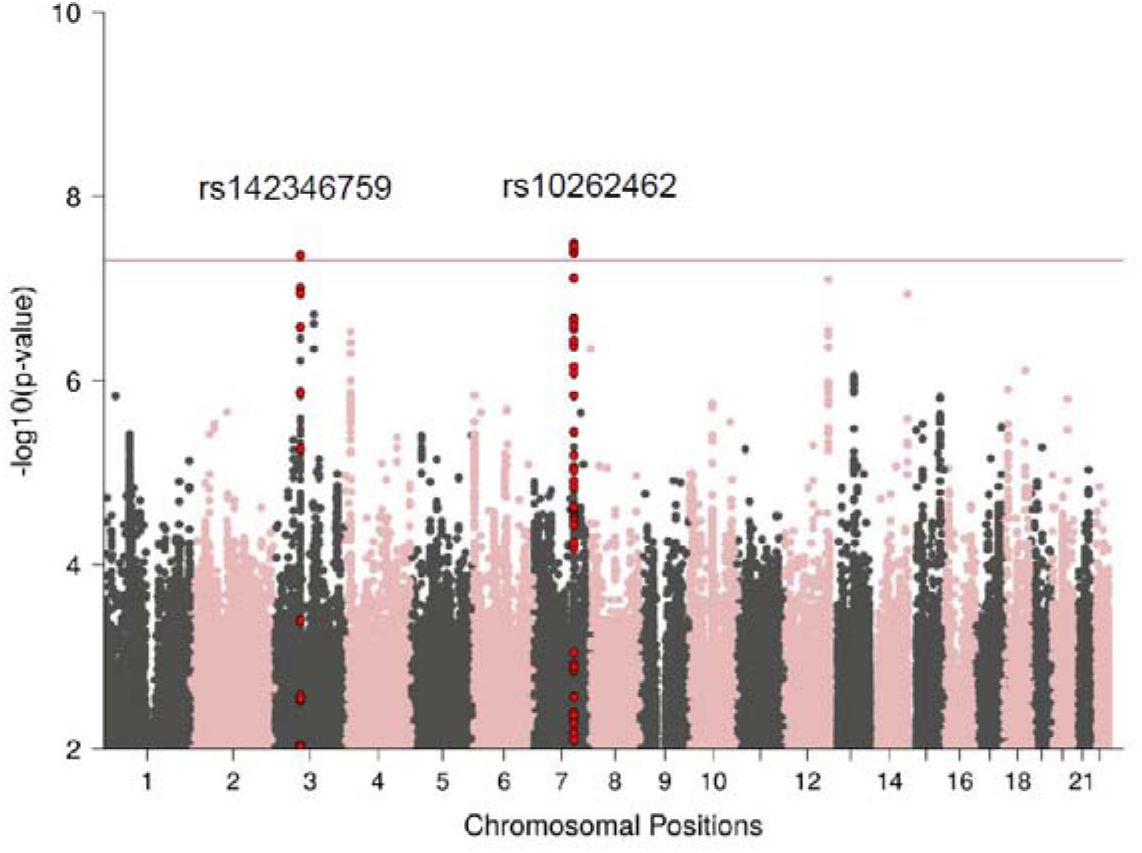
Manhattan plot of UKBB GWAS for childhood maltreatment, showing the top variants. The horizontal line represents genome-wide significance at p<5×10^−8^.

**Figure 2:**
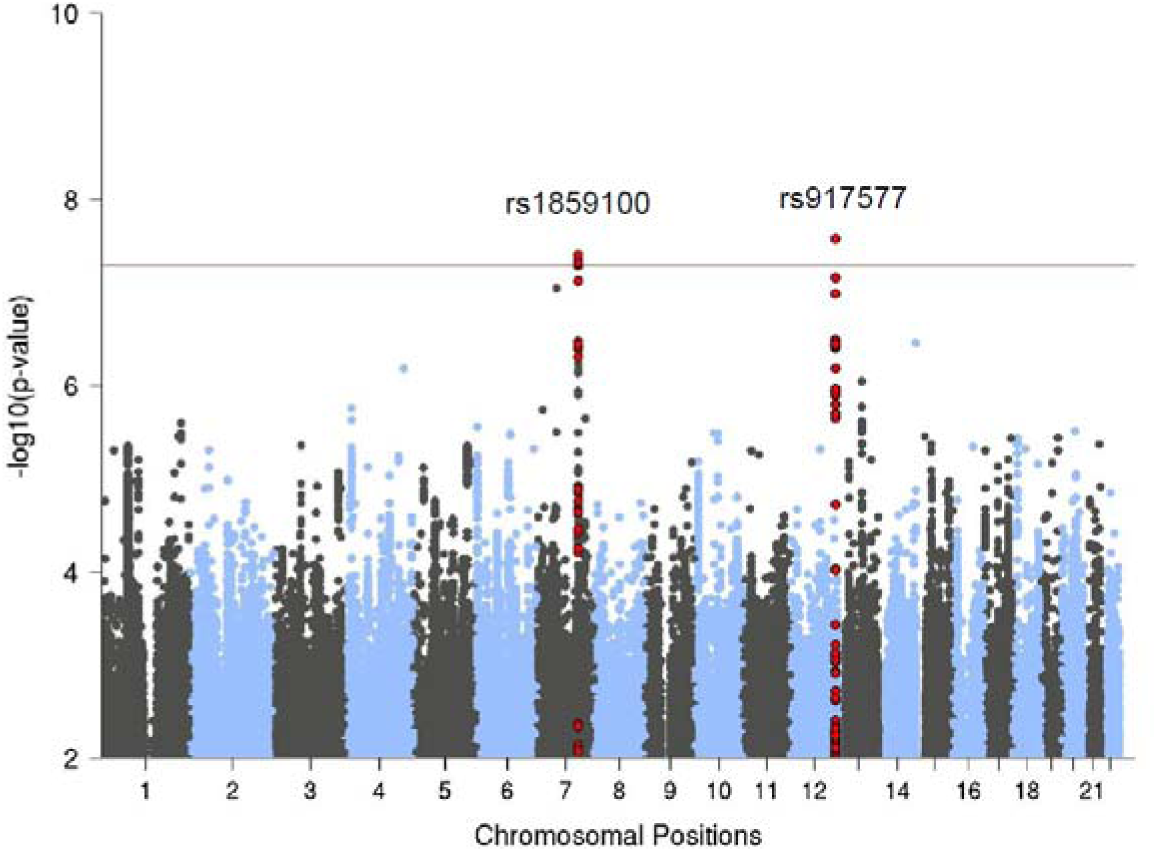
Manhattan plot of the meta-analysis results for childhood maltreatment, showing the top variants. The horizontal line represents genome-wide significance at p<5×10^−8^.

**Figure 3:**
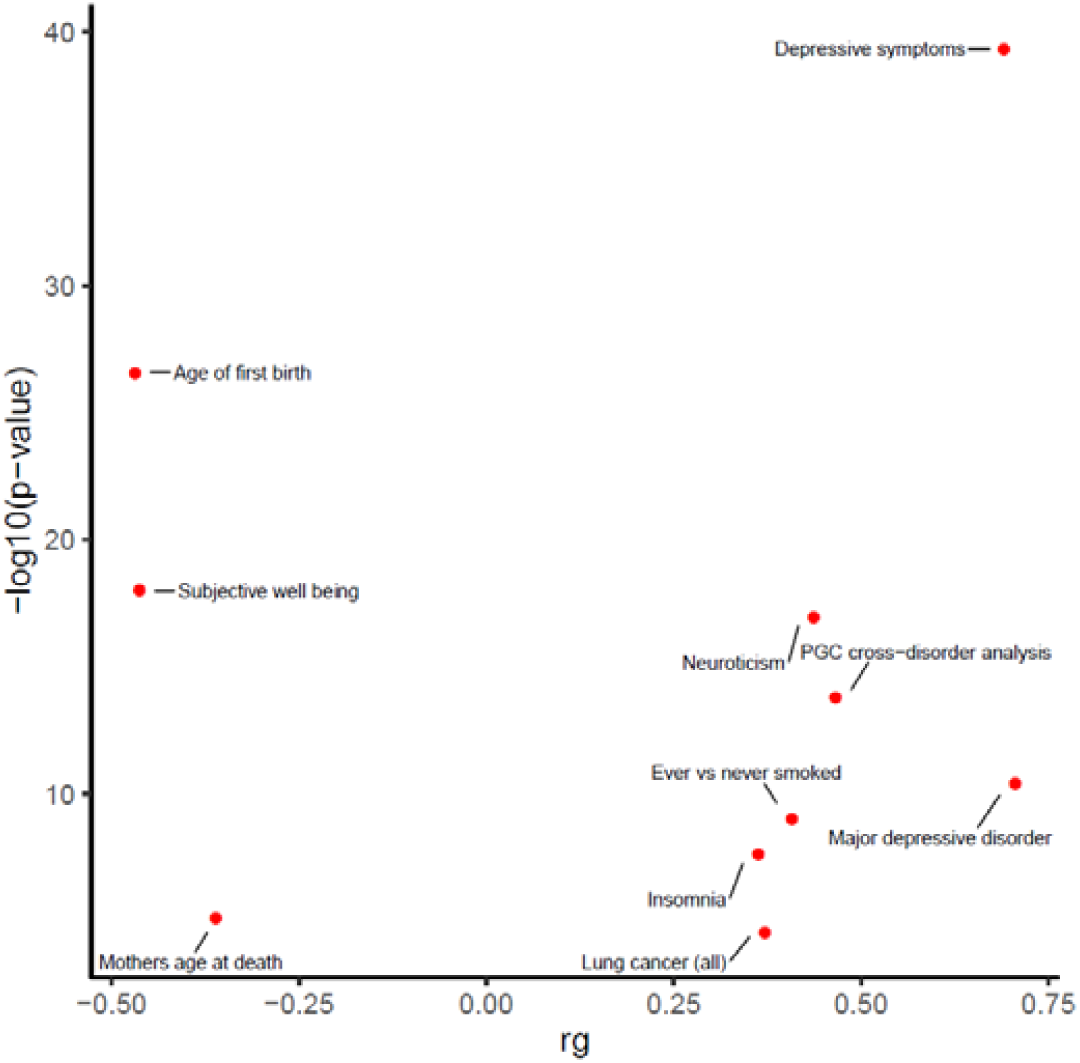
Top ten genetic correlations between several groups of traits (from psychiatric, anthropomorphic, smoking behavior, reproductive, aging, education, autoimmune, and cardio-metabolic categories) and childhood maltreatment (meta-analysis).

Quantile-quantile (qq) plots indicate minimal inflation of p-values across studies (**Supplementary Figures 4-6**). Using the LDSR intercept method, polygenic effects account for 93% and 94% of the observed inflation in test statistics for the UKBB dataset (intercept=1.0096, SE=0.0064) and meta-analysis (intercept=1.0095, SE=0.0077), respectively (**Supplementary Figures 4 and 6**), consistent with minimal population stratification and cryptic relatedness. Heterogeneity estimates in the meta-analyses were not significant (**Table 1**).

### Integration with Functional Genomic Data

Using the web-based tool FUMA, the two UKBB GWAS hits were each annotated to two genes, *FOXP1* and *FOXP2* (**Table 2**). Gene-based analysis of the UKBB GWAS summary statistics further identified three gene-wide significant genes, *KIF26B* (p=1.67×10^−7^), *CNTNAP5* (p=8.89×10^−7^), and *EXOC2* (p=2.04×10^−6^) from a total of 18,989 protein coding genes. Gene-set analysis did not reveal any significant pathways associated with childhood maltreatment. Limited functionality of the two risk variants (rs142346759 and rs10262462) was observed (**Table 2**). One of the SNPs in LD for the risk variant on chromosome 3, rs142346759, obtained a CADD score of greater than 12.37, indicating that this SNP may be deleterious. Six of the SNPs in LD with the risk variant on chromosome 7, rs10262462, had a CADD score of greater than 12.37. No significant eQTLs were identified for either risk locus.

The chromosome 7 variant identified in the meta-analysis, rs1859100, also mapped to the gene *FOXP2* and is located in the same genomic risk locus (chr7:114015707-114287116) as rs10262462. The other hit observed in the meta-analysis, rs917577, was mapped to an intergenic region on chromosome 12. This variant obtained an RDB categorical score of 2B, indicating that it is likely to affect transcription factor binding. No eQTLs exist in the selected tissue types for this region (**Table 2**).

### Heritability of Reported Childhood Maltreatment

GWAS summary statistics were used to estimate the SNP-based heritability (h^2^_snp_) of childhood maltreatment with the tool LDSR (**Table 3**). The h^2^_snp_ was estimated at 0.057 (p=1.60×10^−32^) for the UKBB discovery dataset and 0.123 (p=0.002) for PGC1.5. The h^2^_snp_ for the meta-analysis was 0.057 (p= 4.48×10^−46^).

### Genetic Correlations of Reported Childhood Maltreatment with Other Traits and Disorders

The r_g_ for childhood maltreatment between the UKBB and PGC1.5 datasets was 0.63 (p=3.28×10^−6^). To determine whether there is significant genetic overlap between childhood maltreatment and other traits and disorders, pairwise genetic correlations were calculated using the web-based tool LD Hub. A total of 27 significant correlations (Bonferroni-corrected p-value threshold= 0.05/247=0.0002) were found between childhood maltreatment in the meta-analysis and 247 non-UKBB traits. The top 10 highest genetic correlations are plotted in **Figure 3** with depressive symptoms (r_g_=0.70, p=4.65×10^−40^) having the most significant correlation with childhood maltreatment. There were also positive genetic correlations with “Major depressive disorder” (r_g_=0.71, p=4.13×10^−11^), “PGC cross-disorder analysis” (r_g_=0.47, p=1.62×10^−14^) and “neuroticism” (r_g_=0.44, p=1.14×10^−17^). Significant negative genetic correlations between childhood maltreatment and “age of first birth” (r_g_=-0.47, p=2.61×10^−27^), “subjective well-being” (r_g_=-0.46, p=1.00×10^−18^) and “mother’s age at death” (r_g_=-0.36, p=7.42×10^−6^) were also observed.

## Discussion

The main findings of this study were that 1) variants located in the genes *FOXP1* and *FOXP2* and on chromosome 12 are significantly associated with childhood maltreatment, 2) the SNP-based estimate of childhood maltreatment is approximately 6%, 3) childhood maltreatment is significantly genetically correlated with “depressive symptoms” and “major depressive disorder”, “neuroticism”, “age of first birth”, and “subjective well-being”.

Two genome-wide loci for childhood maltreatment identified in our discovery dataset were also significant in the meta-analysis: rs142346759 (chr3p13), an intronic variant in *FOXP1* and rs10262462 (chr7q31.1) an intronic variant located in *FOXP2*. Both genes form part of the forkhead box superfamily of transcription factors which are widely expressed, and which play important roles during development and adulthood. *FOXP1* and *FOXP2* fall under the FOXP sub-family (also comprising *FOXP3* and *FOXP4*) which has functions in oncogenic and tumour suppressive pathways^38^. *FOXP2* contains highly conserved genomic sites, including an intronic region within this gene, located about 107kb downstream from our risk variant^39^. FOXP1 and FOXP2 have approximately 60% homology at the amino acid level (https://www.ncbi.nlm.nih.gov/books/NBK7023/) and both proteins have been implicated in cognitive disorders, including expressive language impairment^40^. In the meta-analysis, we observed an additional genome-wide variant, located in an intergenic region on chromosome 12, but as this variant does not map to a particular gene, its possible biological mechanism is unclear.

Notably, variation within *FOXP1* has been found to have associations with language impairment, internalizing symptoms, and externalizing symptoms^41^. *FOXP2* has mainly been investigated in regards to speech and language development^42^, but has also been found to be associated with depression^43^ and attention deficit hyperactivity disorder (ADHD)^44^. Further, an intronic variant in the *FOXP2* gene, rs727644, has been associated with risk taking behavior^45,46^. While most work on childhood maltreatment has emphasized subsequent risk for mental and physical disorders, it is possible that externalizing behaviors increase risk for childhood trauma^47^, consistent with a non-passive rGE mechanism. Alternatively, phenotypes such as depression or neuroticism may increase the likelihood of individuals recalling childhood maltreatment^48,49^.

In this study we estimated SNP-based heritability for childhood maltreatment to be ∼6%. A first possibility, in line with a link between *FOXP* variants and externalizing symptoms, is that genetic factors influence environmental factors indirectly through personality and behavior ^9^. A second possibility, consistent with a link of *FOXP* variants with internalizing symptoms and depression, is that genetic factors influence the recall of childhood maltreatment. In particular, retrospective assessment of childhood maltreatment may be limited by recall bias and the respondent’s subjective assessment of the event^50,51^. Indeed, a recent systematic review found very low concordance between prospective and retrospective measures of childhood maltreatment^52^ and those who retrospectively report childhood adversity are at a greater risk for having psychopathology than those who prospectively reported childhood maltreatment^53^.

A twin-based study estimated the heritability of reported childhood maltreatment (comprising physical, and sexual maltreatment and neglect) to be 6%^7^, the same as our SNP-based estimate. As twin-based studies capture latent heritability across the entire genome, these heritability estimates are generally higher than SNP-based heritability estimates, which are limited to common variation and by the number of markers present and tagged on the genotyping array used^15^. However, in this twin study, when considering each maltreatment category separately, the heritability of childhood physical maltreatment, sexual maltreatment, and neglect was 28%, 0%, and 24%, respectively. This suggests that only physical abuse and neglect are heritable and that sexual abuse is not genetically influenced. It is notable that these twin data, then, do not support an rGE for some abuse types.

Our finding of positive genetic correlations between childhood maltreatment, depressive symptoms, and major depressive disorder further suggests that the genetic factors predisposing to reporting early life maltreatment overlap with the genetic variation underlying depression. Genetic correlations between depression, stressful life events and lifetime trauma have led to the hypothesis that genes increasing risk for the development of depression predispose individuals to entering into adverse environments^54,55^. Depressed individuals with and without trauma exposure differ in associated genetic variation, with trauma-exposed individuals having greater SNP-based heritability, supporting this hypothesis^26,56^. On the other hand, polygenic scores for major depressive disorder (MDD) were associated with greater reporting of stressful life events in individuals with MDD^57^. Indeed, current mood can influence the recall of childhood experiences, and individuals with current depression are at an increased likelihood of reporting early life adversity^58^.

In addition to depression, we found significant positive genetic correlations between childhood maltreatment and “neuroticism” and “PGC cross-disorder analysis” (comprised of GWAS summary statistics of five psychiatric disorders: autism spectrum disorder, attention deficit-hyperactivity disorder, bipolar disorder, MDD, and schizophrenia). We observed negative genetic correlations of childhood maltreatment with “age of first birth” and “subjective well-being”. Associations between early life maltreatment and each of these above traits have previously been observed^56, 59-67^. Further investigation is required to delineate the mechanisms that play a role in the relationship between childhood maltreatment and these outcomes.

Our study had a number of limitations. First, the genetic correlation between the UKBB and PGC1.5 datasets was only 0.63, indicating differences between the datasets, which possibly explains the non-replication of our top hit and of greater SNP heritability in PGC1.5. The UKBB dataset comprises healthy volunteers who are typically of a higher socioeconomic status and in better overall health than the general population of comparable age^68^. Thus, the findings reported here may not be generalizable to the general population. However, it is also worth noting that the top hits were significant in the meta-analysis, where additional hits for childhood maltreatment were detected in an intergenic region on chromosome 12.

Second, although many of the study sites included in the final meta-analysis utilized the well-validated CTQ, childhood maltreatment was measured in a diversity of ways for many of the other studies. Thus, our main phenotype was not homogenous and may reflect different aspects of childhood maltreatment across contributing studies. However, we performed GWAS for each study separately, and meta-analysis was conducted with *p-value* and *direction of effect* to minimize heterogeneity. In addition, based on Cochran’s Q-test, none of our significant hits in the meta-analysis showed significant heterogeneity across studies.

This is the first large-scale genetic study to identify specific variants associated with self-reported childhood maltreatment. Variation in *FOXP* genes and other variants associated with childhood maltreatment may put individuals at greater risk for maltreatment. Alternatively, however, these variants may be associated with a greater likelihood of reporting maltreatment, given their association with depression and neuroticism. With the data available, we are unable to indicate definitively which of these explanations is a better one, and it is possible that different mechanisms have more robust explanatory power in accounting for different abuse subtypes as well as different associated psychopathologies. A clearer understanding of the genetic relationships of childhood maltreatment, including particular abuse subtypes, with a range of different psychiatric disorders, may ultimately be useful in developing targeted treatment and prevention strategies.

## Supporting information

Main Results-Tables

## Acknowledgements

This work was funded by Cohen Veterans Bioscience, the NIMH/U.S. Army Medical Research and Materiel Command Grant R01MH106595 to CMN, IL, KJR, and KCK, One Mind, and supported by 5U01MH109539 to the Psychiatric Genomics Consortium. Statistical Analysis were carried out on the NL Genetic Cluster computer (URL) hosted by SURFsara. Genotyping of samples was supported in part through the Stanley Center for Psychiatric Genetics at the Broad Institute of MIT and Harvard. This research has been conducted using the UK biobank resource under application number 16577.

This work would not have been possible without the contributions of the investigators who comprise the PGC-PTSD working group, and especially the more than 206,000 research participants worldwide who shared their life experiences and biological samples with PGC-PTSD investigators. Full acknowledgements are in the **Supplementary Note**.

