## Supplementary-Figures for "Genomic Influences on Self-Reported Childhood Maltreatment"

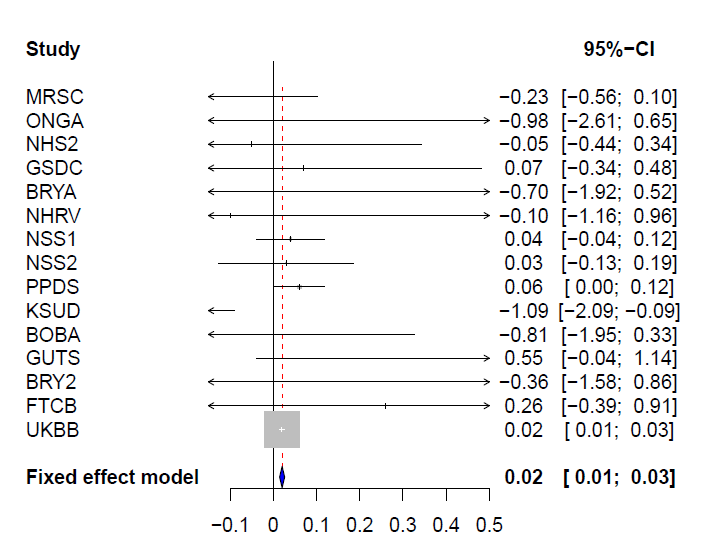

Supplementary Figure 1: Forest plot of the hit rs142346759 (chr3), showing effect sizes for each of the included GWAS in the meta-analysis.

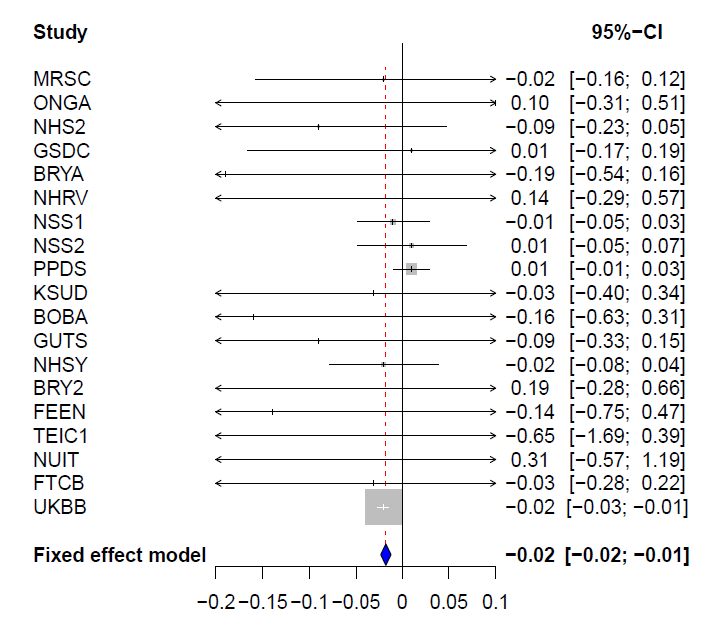

Supplementary Figure 2: Forest plot of the hit rs10262462 (chr7), showing effect sizes for each of the included GWAS in the meta-analysis.

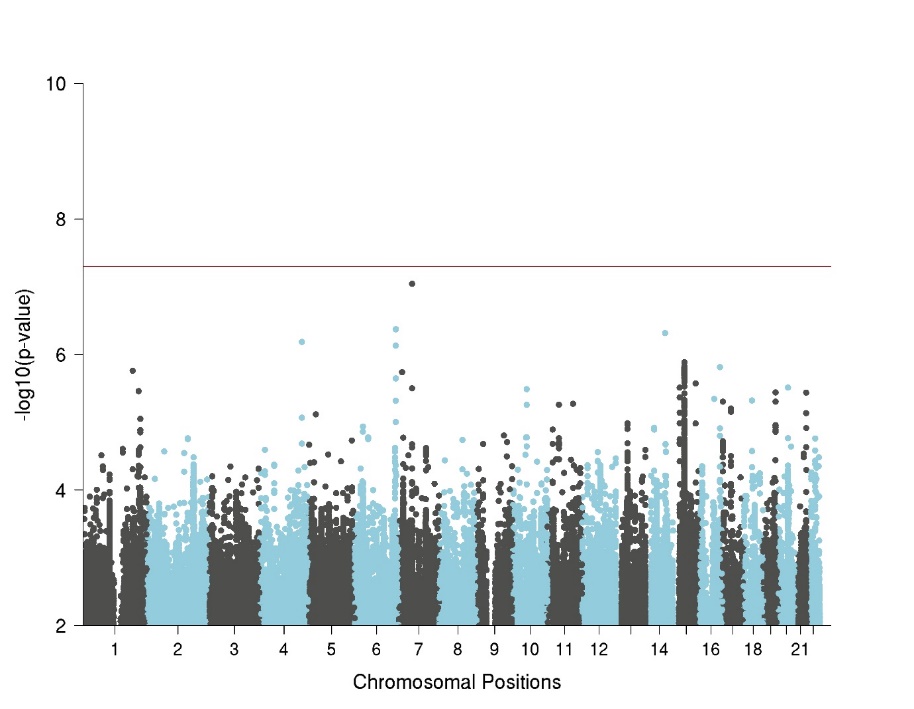

Supplementary Figure 3: Manhattan plot of PGC1.5 GWAS for childhood maltreatment. The horizontal line represents genome-wide significance at p<5x10^-8^.

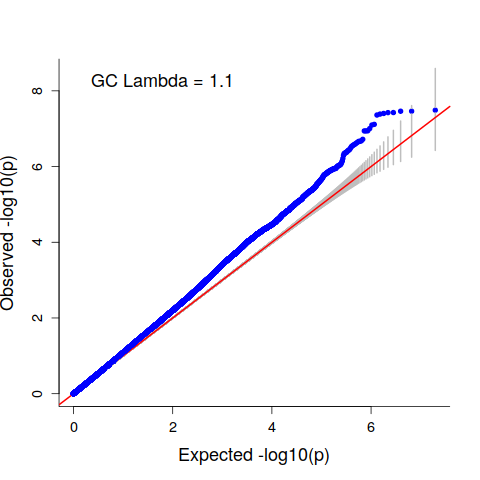

Supplementary Figure 4: Quantile-quantile (QQ) plots of expected versus observed -log_10_ p-values for genome-wide association studies (GWAS) for childhood maltreatment in the UKBB.

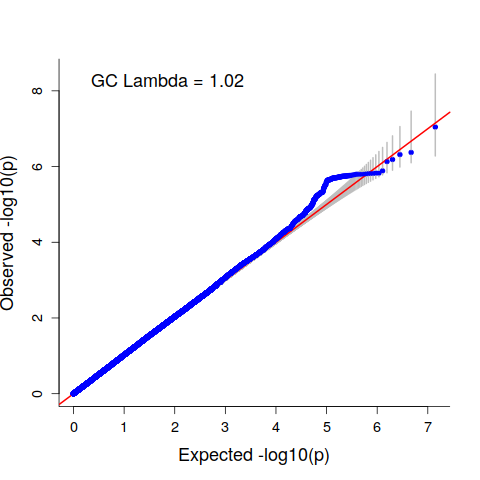

Supplementary Figure 5: Quantile-quantile (QQ) plots of expected versus observed -log_10_ p-values for PGC1.5 results.

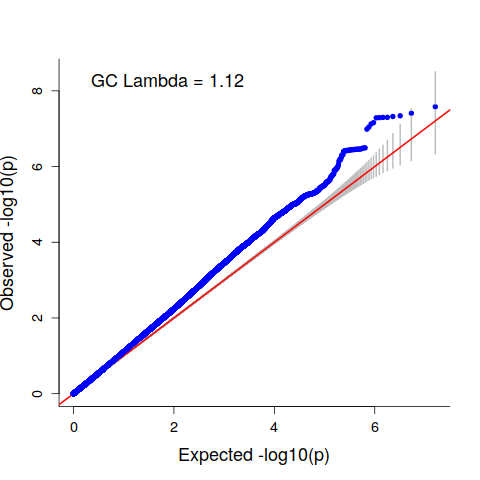

Supplementary Figure 6: Quantile-quantile (QQ) plots of expected versus observed -log_10_ p-values for the meta-analysis results.
