## Supplementary Note for "Genomic Influences on Self-Reported Childhood Maltreatment"

**Study specific acknowledgements**

**Marine Resiliency Study (MRSC)(T1#1)**
The Marine Corps, Navy Bureau of Medicine and Surgery (BUMED) and VA Health Research and Development (HSR&D) provided funding for MRS data collection and analysis (PI DGB) and NIH R01MH093500 funded the GWAS assays and analysis (PI CMN). Acknowledged are Victoria B. Risbrough Ph.D (VA San Diego Healthcare System & UCSD), Mark A. Geyer (UCSD), Daniel T. O’Connor (UCSD), all MRS investigators, as well as the MRS administrative core and data collection staff listed in the Methods article (Baker et al, Prev Chronic Dis. 2012;9(10):E97). The authors also thank the Marine and Navy Corpsmen volunteers for their military service and participation in MRS.

**Ohio National Guard (ONGA)(T1#2)**
The funding information for the Ohio Army National Guard cohort is:
Dept. of Army, Telecommunication and Advanced Technology Research Center (TATRC) Award #s W81XWH-15-1-0080 and W81XWH-10-1-0579 “Ohio Army National Guard Mental Health Initiative: Genetics of Risk and Resilience for Deployment-Related Stress Disorders”.

**Nurses Health Study II (NHS2, NHSY)(T1#5)(T1#22)**
NHSII PTSD Sub-Study was funded by National Institute on Mental Health awards RO1 MH093612, MH078928 to Karestan C Koenen. The NHSII cohort is funded in part by UM1 CA176726.

**Yale-Penn Study (GSDC)(T1#6)**
This study was supported by National Institutes of Health Grants RC2 DA028909, R01 DA12690, R01 DA12849, R01 DA18432, R01 AA11330, and R01 AA017535 and the Veterans Affairs VISN 1 and VISN 4 Mental Illness Research, Educational, and Clinical Centers; and the VA National Center for PTSD Research.
Genotyping services for a part of our genome-wide association study were provided by the Center for Inherited Disease Research and the Yale Center for Genome Analysis. Center for Inherited Disease Research is fully funded through a Federal contract from the National Institutes of Health to The Johns Hopkins University (contract number N01-HG-65403).

**Ash Wednesday and IVS (BRYA)(T1#10)**
This project was funded by a National Health and Medical Research Council Grant (1073041).

**National Health and Resilience in Veterans Study (NHRV; Supplementary Table 1 #13)**
The National Health and Resilience in Veterans Study is supported by the U.S. Department of Veterans Affairs National Center for Posttraumatic Stress Disorder.

**Army Study to Assess Risk and Resilience in Servicemembers (NSS1, NSS2, PPDS)(T1#14, T1#15, T1#16)**
Funding:
Army STARRS was sponsored by the Department of the Army and funded under cooperative agreement number U01MH087981 (2009-2015) with the National Institutes of Health, National Institute of Mental Health (NIH/NIMH). Subsequently, STARRS-LS was sponsored and funded by the Department of Defense (USUHS grant number HU0001-15-2-0004). The contents are solely the responsibility of the authors and do not necessarily represent the views of the Department of Health and Human Services, NIMH, the Department of the Army, or the Department of Defense.

The Army STARRS Team consists of:
Co-Principal Investigators: Robert J. Ursano, MD (Uniformed Services University of the Health Sciences) and Murray B. Stein, MD, MPH (University of California San Diego and VA San Diego Healthcare System)
Site Principal Investigators: Steven Heeringa, PhD (University of Michigan), James Wagner, PhD (University of Michigan) and Ronald C. Kessler, PhD (Harvard Medical School)
Army liaison/consultant: Kenneth Cox, MD, MPH (US Army Public Health Center)
Other team members: Pablo A. Aliaga, MS (Uniformed Services University of the Health Sciences); COL David M. Benedek, MD (Uniformed Services University of the Health Sciences); Susan Borja, PhD (NIMH); Tianxi Cai, ScD (Harvard School of Public Health); Laura Campbell-Sills, PhD (University of California San Diego); Chia-Yen Chen, ScD (Harvard Medical School); Carol S. Fullerton, PhD (Uniformed Services University of the Health Sciences); Nancy Gebler, MA (University of Michigan); Joel Gelernter, MD (Yale University); Robert K. Gifford, PhD (Uniformed Services University of the Health Sciences); Feng He, MS (University of California San Diego); Paul E. Hurwitz, MPH (Uniformed Services University of the Health Sciences); Sonia Jain, PhD (University of California San Diego); Kevin Jensen, PhD (Yale University); Kristen Jepsen, PhD (University of California San Diego); Tzu-Cheg Kao, PhD (Uniformed Services University of the Health Sciences); Lisa Lewandowski-Romps, PhD (University of Michigan); Holly Herberman Mash, PhD (Uniformed Services University of the Health Sciences); James E. McCarroll, PhD, MPH (Uniformed Services University of the Health Sciences); Adam X. Maihofer (University of California San Diego); Colter Mitchell, PhD (University of Michigan); James A. Naifeh, PhD (Uniformed Services University of the Health Sciences); Tsz Hin Hinz Ng, MPH (Uniformed Services University of the Health Sciences); Caroline M. Nievergelt, PhD (University of California San Diego); Matthew K. Nock, PhD (Harvard University); Stephan Ripke, MD (Harvard Medical School); Nancy A. Sampson, BA (Harvard Medical School); CDR Patcho Santiago, MD, MPH (Uniformed Services University of the Health Sciences); Ronen Segman, MD (Hadassah University Hospital, Israel); Jordan W. Smoller, MD, ScD (Harvard Medical School); Xiaoying Sun, MS (University of California San Diego); Erin Ware PhD (University of Michigan); LTC Gary H. Wynn, MD (Uniformed Services University of the Health Sciences); Alan M. Zaslavsky, PhD (Harvard Medical School); and Lei Zhang, MD (Uniformed Services University of the Health Sciences).

**Bounce Back Now (BOBA)(T1#18)**
This work was supported by 1R01MH081056 (PI: Ruggiero), 1R01MH081056-S1 (PI: Amstadter), as well as K02 AA023239 (PI: Amstadter) and K01 AA025692 (PI: Sheerin).

**Cohen Veterans Center Study and Fort Campbell study (COM1, FTCB)(T1#50, T1#52)**
These studies were supported by the Steve and Alexandra Cohen Foundation, Cohen Veterans Bioscience, and the Department of Defense (DoD: W81XWH-09-2-0044 to C.R.M. and W911NF-09-1-0298 to R.Y.).

**Sydney Neuroimaging (BRY2)(T1#36)**
This project was funded by a National Health and Medical Research Council Grant (1073041).

**OPT and CHOICE (FEEN)(T1 #37)**
This research is funded by the National Institute of Mental Health (NIMH; R01MH066347, R01MH066348) and the William T. Dahms, MD, Clinical Research Unit, funded under the Cleveland Clinical and Translational Science Award (UL1 RR024989).
Pfizer Inc. supplied the medication at no cost but had no input in the trial development, conduct, analysis, or interpretation. Drs. Zoellner, Roy-Byrne, Mavissakalian, and Feeny have no competing interests to disclose.
We would like to thank all participants, therapists, and psychiatrists involved in the study and acknowledge the vital contributions of study researchers and administrators in Seattle, Washington and Cleveland, Ohio. Specifically, we would like to thank the investigative team on the grants: Jason Doctor, Ph.D., Joshua McDavid, MD, Alice S. Friedman, MSN, ARNP, and Nora McNamara, MD. Afsoon Eftekhari, Ph.D., and Lisa Stines Doane, Ph.D. were integral in the implementation of this study. Edna Foa, Ph.D. and her team provided PE integrity ratings. We would like to acknowledge Susan Silva, Ph.D., Eric Youngstrom, Ph.D., Kevin King, Ph.D., and Andrew A. Cooper, Ph.D. for their statistical consultation and analyses.

**McLean Trauma Sample (TEIC)(T1#39)**
The Kaufman lab would like to thank the participants for making this research possible; the staff of the Hill Center for Women and Proctor House II, McLean Hospital; The work was supported by National Institute of Mental Health (NIMH) grant R21MH112956 to MLK, and NIMH fellowship grant F32MH109274 to LAML, the Anonymous Women’s Health Fund to MLK, the O’Keefe Family Foundation to MLK, the Trauma Scholars Fund to MLK, and the Frazier Foundation Grant for Mood and Anxiety Research to KJR.
The Teicher Lab would like to thank all the participants for being a part of these studies and the staff of Developmental Biopsychiatry Research Program, McLean Hospital. The studies were supported by National Institute of Mental Health (NIMH) grant RO1 MH91391 and National Institute on Drug Abuse R01 DA17846 to MHT.

**NIU Trauma Orcutt (NIUT)(T1#40)**
This study was supported by the Joyce Foundation (Dr Orcutt) and NIH, HD049907and MH085436.

**UK Biobank (UKBB)(T1#60)**
This research has been conducted using the UK Biobank Resource, as an approved extension to application 16577 (Dr. Breen). This study represents independent research part funded by the National Institute for Health Research (NIHR) Biomedical Research Centre at South London and Maudsley NHS Foundation Trust and King’s College London. The views expressed are those of the authors and not necessarily those of the NHS, the NIHR or the Department of Health and Social Care. High performance computing facilities were funded with capital equipment grants from the GSTT Charity (TR130505) and Maudsley Charity (980). GB and JRIC acknowledge funding from Cohen Veterans Bioscience.
